# Individual behavioral type captured by a Bayesian model comparison of cap making by sponge crabs

**DOI:** 10.1101/330787

**Authors:** Keita Harada, Naoki Hayashi, Katsushi Kagaya

**Author notes:** Corresponding author: Katsushi Kagaya.

## Abstract

‘Animal personality’ is considered to be developed through complex interactions of an individual with its surrounding environment. How can we quantify the ‘personality’ of an individual? Quantifying intra- and inter-individual variability of behavior, or individual behavioral type, appears to be a prerequisite in the study of animal personality. We propose a statistical method from a predictive point of view to measure the appropriateness of our assumption of ‘individual’ behavior in repeatedly measured behavioral data from several individuals. For a model case, we studied the sponge crab *Lauridromia dehaani* known to make and carry a ‘cap’ from a natural sponge for camouflage. Because a cap is most likely to be rebuilt and replaced repeatedly, we hypothesized that each individual crab would grow a unique behavioral type and it would be observed under an experimentally controlled environmental condition. To test the hypothesis, we conducted behavioral experiments and employed a new Bayesian model-based comparison method to examine whether crabs have individual behavioral types in the cap making behavior. Crabs were given behavioral choices by using artificial sponges of three different sizes. We modeled the choice of sponges, size of the trimmed part of a cap, size of the cavity of a cap, and the latency to produce a cap, as random variables in 26 models, including hierarchical models specifying the behavioral types. In addition, we calculated the marginal-level widely applicable information criterion (mWAIC) values for hierarchical models to evaluate and compared them with the non-hierarchical models from the predictive point of view. As a result, the crabs of less than about 9 cm in size were found to make caps from the sponges. The body size explained the behavioral variables namely, choice, trimmed cap characteristics, and cavity size, but not latency. Furthermore, we captured the behavioral type as a probabilistic distribution structure of the behavioral data by comparing WAIC. Our statistical approach is not limited to behavioral data but is also applicable to physiological or morphological data when examining whether some group structure exists behind fluctuating empirical data.

## INTRODUCTION

An individual is an important hierarchical structure in biology. We aim to capture intra- and inter-individual variations in behavior as a probabilistic distribution structure, because it is a prerequisite for the study of ‘animal personality’ (Sih et al., 2004; Niemelä and Dingemanse, 2018). Because the term ‘individual difference’ sometimes means only inter-individual difference, we use ‘individual behavioral type’ to refer to the two variations. Behavioral ecologists and evolutionary biologists have been interested in the behavioral type because behavior can be a selective trait. At the evolutionary time scale, the distribution structure is very likely to be related to the evolvability of behavior (Kirschner and Gerhart, 1998). At the behavioral time scale, the behavioral type can be caused through complex and dynamic interactions of individual properties such as behavioral plasticity based on physiological processes, with surrounding dynamic environments. A typical interaction can be observed in body extending behaviors such as tool making and usage (e.g., Hunt, 1996; Wang et al., 2014; Matsui and Izawa, 2017; Sonoda et al., 2012). A body extending behavior, which is basically a behavior involving the attachment of non-living things to a body, seems to require at least some information processing to infer a current body size in order to achieve an adaptive extension through complex interactions. Uncertainty in the inference, and accumulation of experiences accompanying the realization of body extensions, might result in the emergence of some behavioral type. Here we examine the hypothesis that individual behavioral types would emerge in the body extending behavior. As an example of the body extending behavior, the sponge crab ‘s behavior of cap making and carrying is experimentally examined and statistically modeled in this study.

To capture the structure, we need repeated measurements and specific statistical modeling considering a hierarchical structure (Niemelä and Dingemanse, 2018). Hierarchical models such as a generalized linear mixed effect model (GLMM) are widely considered appropriate for the repeated data (Zuur et al., 2009; Niemelä and Dingemanse, 2018; Reinhart, 2015). However, empirical data has poorly examined the appropriateness of a hierarchical model relative to a non-hierarchical one such as the generalized linear model (GLM). One well-known statistical measure used in GLM from the predictive point of view is the Akaike Information Criterion (AIC) (Akaike, 1974; Sakamoto et al., 1986). To calculate AIC, the maximum log-likelihood needs to be calculated, but in general, prediction by the maximum likelihood (ML) method is inappropriate for hierarchical models (Watanabe, 2005). This is because a model with hierarchical structures is a statistically non-regular model and the assumptions set in the ML estimation are considered inappropriate (Watanabe, 2005, 2010b, 2018). Then, how much is the degree of inappropriateness? Alternatively, a Bayesian procedure to construct a predictive distribution is known to perform better than the ML method in the hierarchical models in terms of the predictive point of view (Watanabe, 2018). The Bayesian framework can give a unified measure of the appropriateness.

Although the basic Bayesian framework and its mathematical foundation of measuring the predictability of an arbitrary pair of a statistical model and a prior distribution, has been rigorously established (Watanabe, 2010b,a, 2018), there are few applications of the framework to behavioral data containing repeated measurements (Wakita et al., 2020). Specifically, the performance of a predictive distribution can be inferred using the Widely-Applicable Information Criterion (WAIC); it is a measure of the generalization error defined as the extent to which a specified predictive distribution is approximated with respect to an unknown true distribution that generates data (Akaike, 1980; Watanabe, 2018). Furthermore, there are almost no appropriate applications of WAIC to hierarchical models for repeatedly measured data. To construct a predictive distribution using a hierarchical model, we are usually interested in a new observation from a new cluster other than from the clusters that provided the initial observations. Therefore, we need to marginalize the parameters assigned to each cluster when training a model to calculate WAIC in that situation (Watanabe, 2018; Millar, 2018). However, this point does not seem to be recognized well not only in biological communities but also generally in other real-world data analyses.

Therefore, we propose and adopt a Bayesian model comparison framework using WAIC to study a specific individual behavioral type in the body extending behavior of the crab. In previous research, one field study dealt with the preference of dromiid crabs for materials and examined the association between a cap size with a body size (McLay, 1983). Additionally, it is reported that *Cryptodromia hilgendorfi* use caps made from many species of sponges, but they particularly prefer the sponge *Suberites carnosus*, and the crabs make sponge caps twice as large as the carapace area. In previous experimental research, the preference for material size and their suitability for the body size and cap size are scarcely investigated. It is reported that *Dromia personata* mainly uses sponges and ascidians (Bedini et al., 2003), although they could also make caps with paper (Dembowska, 1926). Dembowska (1926) reported that a non-breaking space is used qualitatively and that the cap size made by *Dromia personata* (reported as *D. vulgaris*) using paper is as large as the size of the caps originally carried by the crabs. Because these studies once sampled a body size and a camouflage size for an individual, it is unclear whether there is an individual behavioral type. In addition, it is unknown whether a behavioral type that is conditional on the body size exists in the cap making behavior. Thus, although the crabs in the family *Dromiidae* have been known to make a cap (Guinot and Wicksten, 2015), the behavior of the *Lauridromia dehaani* has not been examined so far.

Accordingly, the lack of experimental data on the cap making behavior of the crab *Dromia personata* and the limitations of the statistical approach, we set four goals to study the individual behavioral type in the body extending behavior: (1) to perform behavioral experiments by sampling behavioral data repeatedly, (2) to formulate an individual behavior type in statistical models, and simultaneously to construct other models assuming no such behavior type, (3) to measure the predictive performance of those models by WAIC, including hierarchical models that assume a particular individual behavior type and compare the findings with those of non-hierarchical models assuming the existence of no such behavior type, (4) to infer a relationship between the behavioral data and the body size by conditioning the behavioral variables by the body size.

## MATERIALS & METHODS

### Animal collection

From December 2015 to April 2017, 40 individuals (21 males, 19 females) of *Lauridromia dehaani* (Brachyura: Dromiidae) (Fig. 1A) were captured using a gill net at the Sakai fishing port, Minabe town, Wakayama, Japan (33° 44 ‘N, 135° 20 ‘E). We conducted behavioral experiments of cap making on 38 individuals (20 males, 18 females) and video recorded the behaviors of 2 individuals (4.30 cm and 7.19 cm of the carapace widths for each) in a tank filled with filtered natural seawater (about 3.4 % of the salinity) at Shirahama Aquarium, Seto Marine Biological Laboratory, Kyoto University (33° 41 ‘N, 135° 20 ‘E), from December 2015 to June 2017. For the behavioral experiments, we successfully sampled 8 individuals repeatedly (3 or 5 times for each). Thus, we only sampled one observation from one of the other 30 individuals. Note that although the sample sizes of the behavioral acts for each individual are different our method is still applicable. Before the experiments, all individuals were retained in the tanks (19.5–23.8 °*C*, light on: 8–17, light off: 17–8) of the aquarium for more than two days for acclimation. We measured their carapace width (cm) (Fig. 1B) as a proxy for the body size, and divided them into three levels depending on whether they lacked any of the fourth and fifth pereiopods: (O) none of the fourth and fifth pereiopods were absent, (1) one of them was absent, (2) both fourth and fifth of each side were absent.

**Figure 1.**
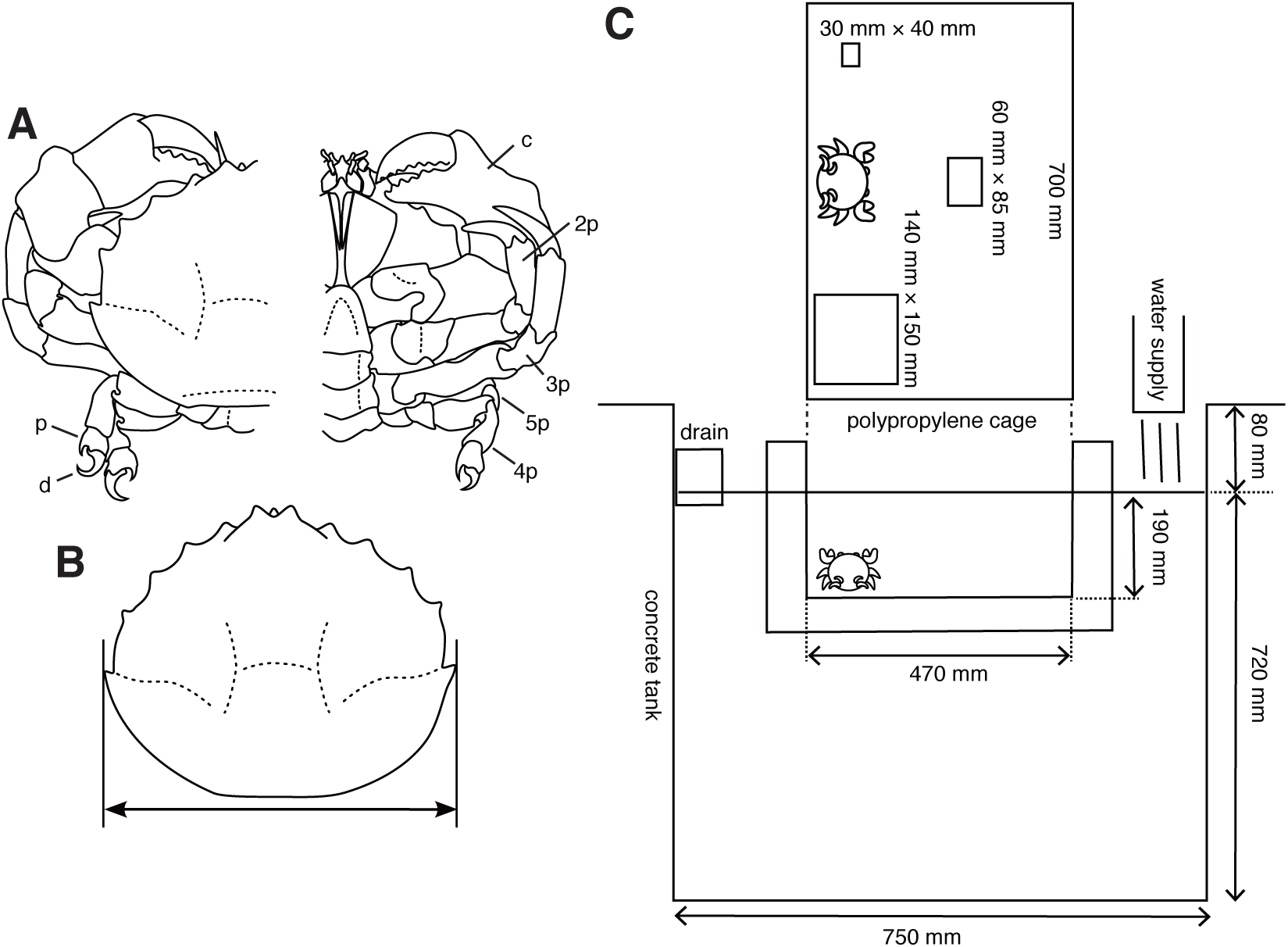
Experimental animals and their setup. (A) *Lauridromia dehaani*; p—propodus of fifth pereiopod; d—dactylus of fifth pereiopod; c—chela (1st pereiopod); 2p—second pereiopod; 3p—third pereiopod; 4p—fourth pereiopod; 5p—fifth pereiopod. (B) Carapace width we measured. (C) Experimental cage floating in an aquarium tank with three different sizes of sponges. The drawings are by the authors.

### Experimental setup and procedure

We prepared three sizes of white melamine foam that is found commonly worldwide (most notably manufactured by BASF of Germany) and often used in general households (in Japan, it is called Gekiochi-kun, LEC, Inc.) (S: 20 mm × 30 mm × 40 mm; M: 30 mm × 60 mm × 85 mm; L: 30 mm × 140 mm × 150 mm). We used this sponge because it is easy to sink.

First, to confirm that the cap making behavior of the crab *Lauridromia dehaani* is similar to the behavior in the reports (Dembowska, 1926; McLay, 1983), we video-recorded the behavior from two crabs. The two crabs were used only for video recording in the aquarium (310 mm × 180 mm × 240 mm, W × L × H). We started the recording from 9 am to 10 am in the morning, and stopped for 2 h after the crabs stopped cap making. We used a red light transmitted through a polyvinyl chloride board and excluded another light source by enclosing the aquarium. We obtained 5 recordings for each crab.

Secondly, we performed experiments on choice of cap size, trimming, and excavating behaviors. S size sponge was smaller than all crabs, whereas the L size was larger than all crabs. Each sponge was put pseudo-randomly on either side and at the back center of the cage (700 mm × 470 mm × 190 mm, W × L × H, Fig. 1C), which floated in the tank. Then, crabs were introduced to the front center of a cage floating in the tank, thereby the distance between each sponge and the crab was equal. We started a trial from 9 am to 10 am and checked whether the crab carried any sponge once a day. We counted the days when the crab carried a sponge. The latency, measured as the number of days to produce a cap, is modeled as a random variable. Note that the crab was assumed to make a cap at night, because it is considered nocturnal (McLay, 1983). If it did, we collected the sponge; otherwise, the crab and the three sponges remained in the cage. When the crab did not carry any sponge for five days, we stopped the trial. We destroyed all sponges that the crabs processed and measured their whole area (*cm*^2^), and area of the concave part (*cm*^2^) excavated by a crab from the pictures taken 46 cm above the sponges. The trimmed area and concave area are modeled as random variables. Although in the beginning we only performed one trial for one individual (*N*_*animal*_ = 30), we obtained five trials for one individual after February 2017 (*N*_*animal*_ = 8) to examine the behavior type. Our hypothesis that a behavioral type would be formed was conceived after the day, but we did not change the experimental condition.

### Statistical modeling

To quantify the behavioral type in the experiment, we constructed 26 statistical models (Table. 1) for the four different aspects: (1) choice of sponge size (6 models), (2) amount of sponge trimmed by cutting (8 models), (3) size of cavity (6 models), and (4) latency to produce a cap (6 models). In each case, we built the statistical models specifying individual behavioral types as hierarchical structures with parameters and performed MCMC samplings from the posterior distribution. Also, we conditioned the variables with the carapace width, levels of leg absence, and gender. We specified the models in the probabilistic programming language Stan (Stan Development Team, 2018). We used non-informative uniform priors for some parameters unless otherwise explicitly described. The performed samplings from the posterior distributions using No-U-Turn Sampler (NUTS) implemented as a Hamiltonian Monte Carlo (HMC) sampler in Stan. Sampling convergence was visually diagnosed by trace plots and quantitatively via the Gelman-Rubin convergence statistic, *R*_*hat*_ (Gelman et al., 1992). All sampled draws were judged to be converged when *R*_*hat*_ < 1.10, were used to construct predictive distributions with WAIC on each model. All computations were performed in the R statistical environment (R Core Team, 2018), and the Stan codes for each model were compiled and executed through the R package *rstan* (Stan Development Team, 2018).

**Table 1.** Summary of model structures and the predictive performances in WAIC. Abbreviations, intercept_L: intercept in the linear predictor (LP) for the choice of L; intercept_1: intercept in the LP for the decision of trimming; intercept 2: intercept in the LP for the mean of the trimmed size of the sponge; CW: carapace width; Leg: degree of the leg lack; _L and _NO: parameters for L sponge and skipping, respectively; Choice: choice of whether to cut the sponge or not; Gender: gender of the animal; intercept_2: intercept in the LP for the mean of the days to carrying; Choice: choice of sponge size; ZIP: Zero-inflated Poisson distribution; WAIC: value of Widely-Applicable Information Criterion per sample; dWAIC: the difference of the WAIC of the model against the best-performed model.

| response variable | model | hierarchical structure | conditioning variables | link function | distribution | WAIC (nat) | dWAIC (nat) | plot |
| --- | --- | --- | --- | --- | --- | --- | --- | --- |
| Choice | 1.1 | intercept_L | CW_L Leg_L CW_NO Leg_NO | softmax | categorical | -2.13 | 0.00 | Fig.3A |
| Choice | 1.2 | intercept_L | CW_L CW_NO | softmax | categorical | -1.87 | 0.26 | – |
| Choice | 1.3 | intercept_L | – | softmax | categorical | -0.88 | 1.25 | – |
| Choice | 1.4 | intercept_L | Leg_L Leg_NO | softmax | categorical | -0.78 | 1.35 | – |
| Choice | 1.5 | – | CW_L CW_NO | softmax | categorical | 0.85 | 2.99 | Fig.3C |
| Choice | 1.6 | – | CW_L Leg_L CW_NO Leg_NO | softmax | categorical | 0.87 | 3.01 | – |
| Trimmed size | 2.1 | intercept_1 intercept_2 | CW Choice | logit log | ZIP | -2.08 | 0.00 | Fig.4A |
| Trimmed size | 2.2 | intercept_2 | Choice | logit log | ZIP | 0.81 | 2.89 | – |
| Trimmed size | 2.3 | intercept_2 | CW Choice | logit log | ZIP | 0.86 | 2.95 | – |
| Trimmed size | 2.4 | intercept_2 | – | logit log | ZIP | 1.23 | 3.32 | – |
| Trimmed size | 2.5 | intercept_2 | CW | logit log | ZIP | 1.37 | 3.46 | – |
| Trimmed size | 2.6 | – | CW Choice | logit log | ZIP | 7.40 | 9.48 | Fig.4B |
| Trimmed size | 2.7 | – | CW | logit log | ZIP | 10.05 | 12.13 | – |
| Trimmed size | 2.8 | – | – | logit log | ZIP | 12.55 | 14.63 | – |
| Cap cavity size | 3.1 | intercept | CW | log | gamma | 4.45 | 0.00 | Fig.5A |
| Cap cavity size | 3.2 | – | CW | log | gamma | 4.54 | 0.08 | Fig.5B |
| Cap cavity size | 3.3 | – | CW Gender | log | gamma | 4.69 | 0.24 | – |
| Cap cavity size | 3.4 | intercept | – | log | gamma | 4.71 | 0.26 | – |
| Cap cavity size | 3.5 | – | CW | identity | normal | 4.75 | 0.30 | – |
| Cap cavity size | 3.6 | intercept cw | CW | log | gamma | 6.18 | 1.73 | – |
| Latency for making | 4.1 | intercept_2 | CW | logit log | ZIP | 1.10 | 0.00 | Fig.6A |
| Latency for making | 4.2 | intercept_2 | – | logit log | ZIP | 1.28 | 0.18 | – |
| Latency for making | 4.3 | – | – | logit log | ZIP | 1.28 | 0.19 | Fig.6B |
| Latency for making | 4.4 | – | Choice | logit log | ZIP | 1.30 | 0.20 | – |
| Latency for making | 4.5 | – | CW | logit log | ZIP | 1.38 | 0.28 | – |
| Latency for making | 4.6 | – | CW Choice | logit log | ZIP | 1.72 | 0.62 | – |

We compared the predictive performances of the models using WAIC (Watanabe, 2018, 2010b). It should be emphasized that the WAIC of a hierarchical model can be defined in several ways depending on how a predictive distribution is defined. In our case, as we would like to construct a new distribution regarding a new act of a new individual, we have to marginalize the intermediate parameters assigned to each individual in the statistical model (Watanabe, 2018). This is because we are interested in the prediction of a new behavioral act when we get a new individual and a new behavioral act instead of the prediction of a new behavioral act from the individuals sampled in this study. By performing this procedure, we can equally compare a hierarchical model with a non-hierarchical model, because the focus of the prediction in a non-hierarchical model is on a new behavioral act of a new individual.

Here we briefly describe the basic procedure based on Watanabe (2018). Let *X*^*n*^ = (*X*_1_, …, *X*_*n*_) be an sample from the unknown true distribution and *p*(*x*|*w*) a statistical model with *w* assigned to each individual. Furthermore, *w* is assumed to be taken from *φ*(*w*|*w*_0_) to form a hierarchical structure. In learning step, 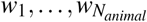 is prepared. In prediction step, our statistical model is built like *p*_*model*_ (*x*|*w*_0_) by marginalizing *w* out:

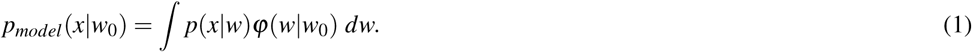

WAIC is a measure for the degree of accuracy of an approximation of a predictive distribution to the true distribution generating data. For our hierarchical model, the predictive distribution is defined as 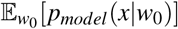. Then, the marginal-level WAIC for a hierarchical model is defined as:

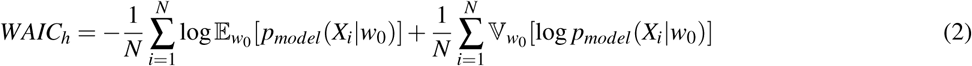

where 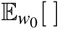 and 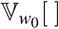 are the average and variance operators, respectively, of the posterior distribution of *w*_0_. *w*_0_ is estimated so that practically the MCMC sample is used, thus numerical integration is required. In this study, the computation is implemented in the ‘function’ block in the Stan codes using the Simpson’s rule for the numerical integration and the *log_sum_exp* function provided in Stan (see supplementary material).

On the contrary, the WAIC for a non-hierarchical model is defined for a statistical model *p*_*model*_ (*x*|*w*):

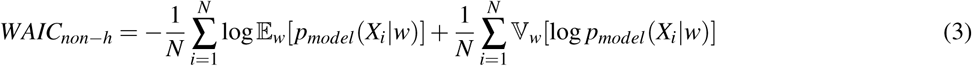

where 𝔼_*w*_[ ] and 𝕍_*w*_[ ] are the average and variance operators, respectively, of the posterior distribution of *w*. Note that the often used conditional-level WAIC is described in the Discussion.

#### Choice of material size (model 1_1)

To provide an overview of the specified models, we describe only the best-performing models in terms of WAIC here. The other models are summarized in Table 1.

We formulate a tendency toward a choice as *µ*[*n, m*] (*m* = 1, 2, 3 for M, L, skip, respectively):

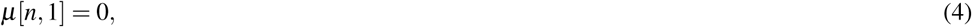

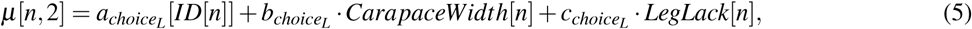

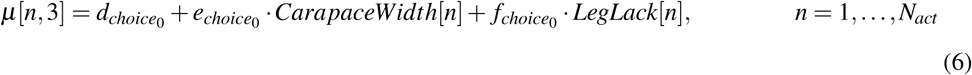

where *N*_*act*_ = 68 is the total number of behavioral acts, and *ID* represents animal identity (from 1 to *N*_*animal*_ =38). *µ* is linked to the linear predictor in terms of the carapace width, *CarapaceWidth* and the level of absence of leg, *LegLack*. The choice of an M size is fixed to zero. 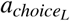 is for each individual, thus it is hierarchized. 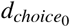 is not hierarchized. The distribution of 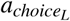 is defined as the normal distribution with the mean 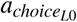 and standard deviation 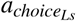:

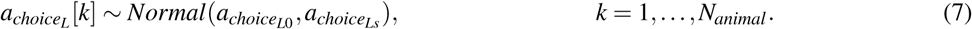

The actual choice *Choice* is defined as the categorical distribution with the softmax function:

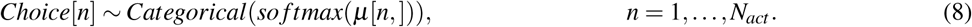

Thus, in this case, a statistical model *p*_*model*_ (*x*|*w*_0_) is set using the parameters:

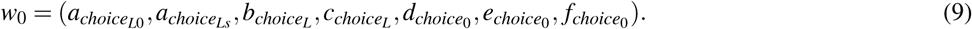

Note that 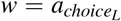 is marginalized out when we build the predictive distribution, so that it is not included in *w*_0_. The choice *Choice*[*n*] is modeled as a random variable *X*_*n*_. *CarapaceWidth*[*n*] and *LegLack*[*n*] are the conditioning variables.

#### Trimming (model 2_1)

The probability of making the decision on whether an animal is cut off the sponge is written as *ϕ*_*cut*_ linked to the linear predictor with the carapace width *CarapaceWidth* and the selected sponge size *Choice*[*n*]:

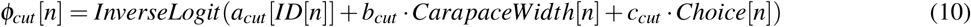

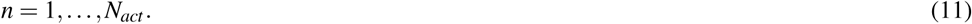

*a*_*cut*_ is assigned for each individual. *N*_*act*_ = 51 and *ID* is from 1 to *N*_*animal*_ = 30. The distribution of *a*_*cut*_ is defined as the normal distribution with the mean 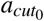 and standard deviation 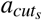:

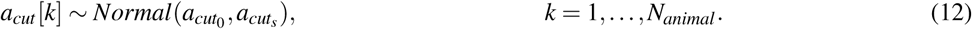

The prior distribution of 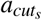 is defined as the half *t* distribution:

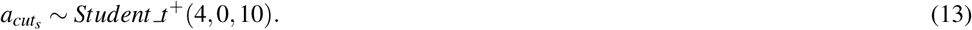

The mean area of a sponge trimmed by the crab *λ* is linked to the linear predictor with the log link function:

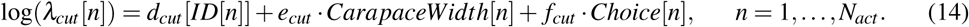

*d*_*cut*_ is assigned for each individual. The distribution of *d*_*cut*_ is defined as the normal distribution with the mean 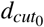 and the standard deviation 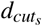:

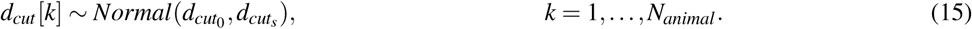

The prior distribution of 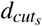 is defined as the half t distribution:

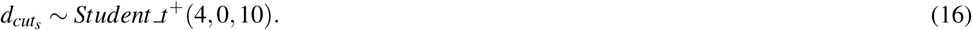

Altogether, the area of the trimmed sponge is modeled as the variable *Trimmed*. Its distribution of it is defined as the zero-inflated Poisson distribution (ZIP) with the parameters *ϕ*_*cut*_ and *λ*_*cut*_:

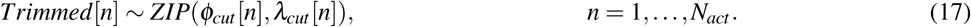

When a crab skips the trimming behavior, *Trimmed* is set to zero even if the sponge size is smaller than the defined M and L sizes owing to measurement errors. Note that *Trimmed* is rounded off to an integer. We assume that the rounding process has no significant impact on the data distribution.

#### Cap cavity making (model 3_1)

To examine the relationship between the cap cavity size *CavitySize* and the carapace width *CarapaceWidth*, gamma distribution is chosen to represent non-negative values of the cavity size. The mean of the distribution is specified by *lambda*_*cavity*_ with shape and rate parameters:

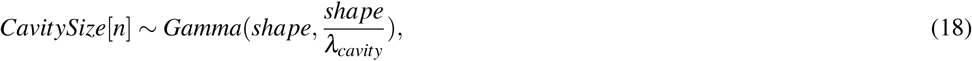

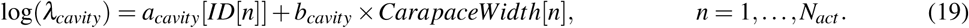

where the rate parameter was given as the shape over the log-linked linear predictor and *a*_*cavity*_ is the intercept for each individual. *N*_*animal*_ = 30, *andN*_*act*_ = 51. The *a*_*cavity*_ is taken from the normal distribution with the mean 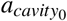 and the standard deviation 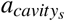:

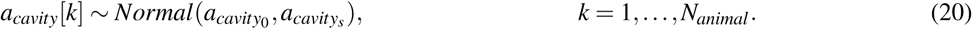

#### latency (model 4_1)

We assume that the latency to produce a cap, *Days*, fits the ZIP distribution, which is similar to the *Trimmed* case:

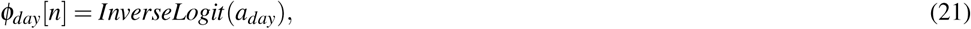

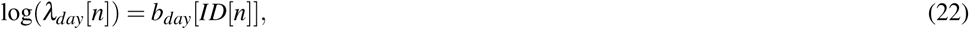

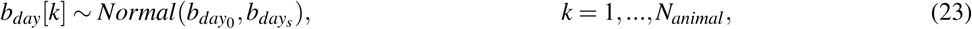

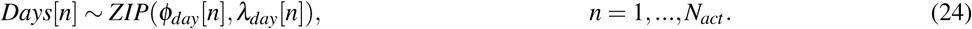

where *N*_*animal*_ = 32, *N*_*act*_ = 56. Note that *b*_*day*_ is into this model to construct a hierarchical structure. All the data and codes are available from the supplementary material.

## RESULTS

We measured and modeled the four variables: the choice of sponge size, trimmed size, cavity size, and latency for completing making sponge, as random variables. Furthermore, we evaluated the model predictability by WAIC (see Materials and Methods).

### Cap making using an artificial sponge

The behaviors of the two crabs were video recorded to confirm the Cap making behavioral sequence when using an artificial sponge (Supplementary Fig. 1). The crabs grasped either side of the sponge using their second and third pereiopods, and trimmed small pieces of the sponge using their chelae (Fig. 2A upper left, upper right, Supplementary movie 1). They first visited the two sides of the sponge. To make a cavity, the crabs rotated their bodies backward and grasped the sponge by the fourth and fifth pereiopods. By repeating these behaviors, the crabs made a groove to cut off a portion of a sponge. In 9 of the 10 trials conducted, it took about 50 min to cut the portion, and the crabs started excavating as soon as they finished the trimming behavior. In the other trial, it took 19 min. Next, the crabs made cavities by tearing off small pieces of a sponge (Fig. 2A bottom, Supplementary movie 2). It took 11 min on average to excavate the cavity. Then, the crabs rotated their bodies backward in order to catch the excavated sponges with the fourth and fifth pereiopods while they kept the portion grasped by the second and third pereiopods. Finally, the crabs released the second and third pereiopods from the cap and carried it off (Fig. 2B, C). In terms of behavior, the crabs often rotated their bodies forward, dorso-ventrally, to enlarge the cavity. It is rare for them to move laterally. They repeated the excavation activities up to eleven times per night and it took up to 4 h. When the crabs rotated their bodies, the direction of rotation was maintained along with the sponge. While the crabs cut the sponge, they actively moved around the sponge. In contrast, they persistently stayed under the sponge during excavation.

**Figure 2.**
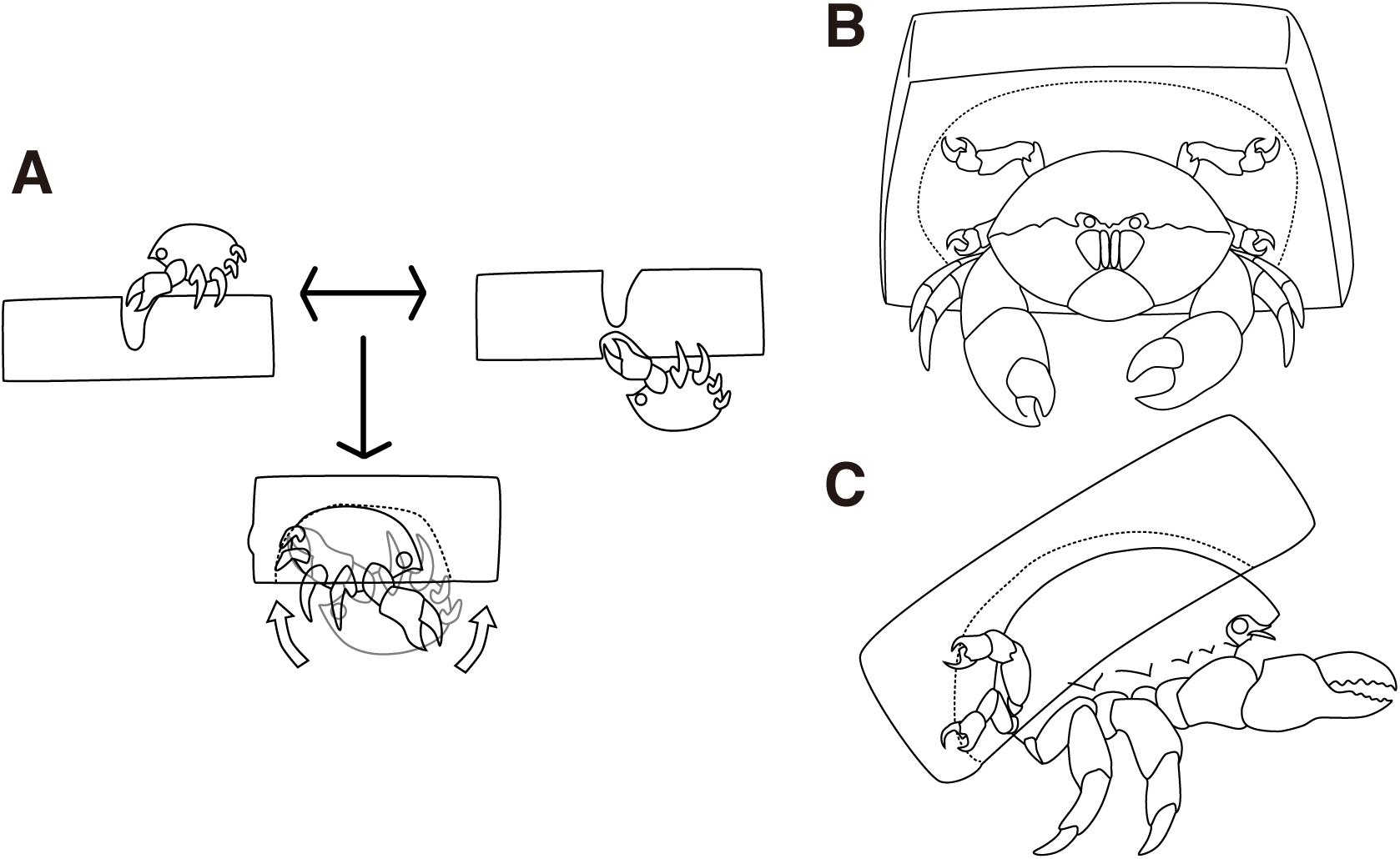
Cap making and carrying behavior. (A) Cap making behavior. (B–C) Carrying behavior of a crab. The drawing is by the authors.

### Sponge choice

None of the 38 animals chose the S size sponge, and 7 animals skipped the cap making behavior (Fig. 3A). Therefore, we defined the choice as a random variable taking either of the three values: M, L, or skipping. The hierarchical model assuming behavioral types 1_1 (Fig. 3A, B) outperformed the non-hierarchical one in terms of WAIC (2.99 nat/sample in the difference, Fig. 3A-D, Table 1). The posterior probability of the behavioral choices was more widely variable on model 1_1 than in model 1_6 depending on the individual difference specified as 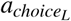 (Fig. 3B). To show the variability, the probability of choice sampled from the posterior distribution from the highest density is visualized in white lines (Fig. 3A,C). Note that the variability of the choice probability in the white curved lines is smaller than the model 1_1 even if the number of lines are the same. Although the body size of the animal indicated with the white arrowhead (Fig. 3A) is small, it preferably selected the size L. This indicates a large inter-individual difference. In the case of either the hierarchical or the non-hierarchical model, the behavioral choice of the sponges was better explained by the carapace width (Fig. 3A, C; Table 1). The estimated information gained by the model 1_1 against model 1_3 is 1.35 nat/sample (Table 1). This suggests that larger crabs tend to choose L size sponge rather than M size. However, the crabs larger than about 9 cm carapace width did not choose the sponges.

**Figure 3.**
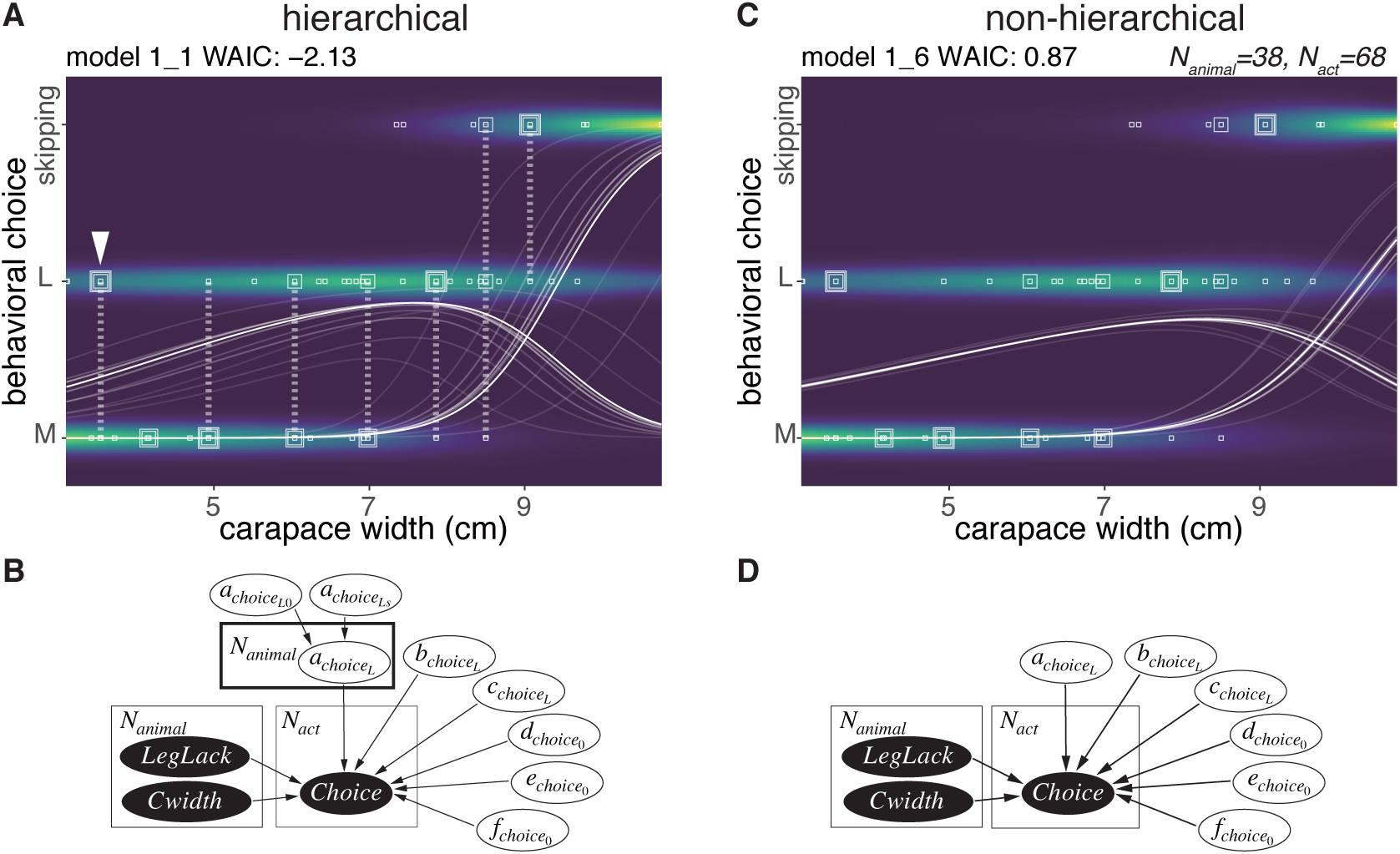
Sponge choice. (A) Predictive distribution on the hierarchical model 1_1 with data points of the behavioral choices, which are M or L size choices or skipping the behavior. The points connected by dotted lines represent data from the same individual. The white curved lines are ten samples from the posterior distribution in decreasing order from the highest density of a parameter representing the probability of a choice. (B) Structure of the model 1_1 in a graphical diagram. 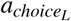 is a parameter assigned to each individual. The variables in the black and white ellipses represent observed data and parameters to be estimated, respectively. (C) Predictive distribution of a choice on the non-hierarchical model 1_6. (D) Structure of the model 1_6 in a graphical diagram.

### Trimming

After a choice of either the M or L size sponges, the crabs decided whether to trim the extra parts of the sponges (Fig. 4A-C). Here, we modeled the size of an area in a sponge that was trimmed (*N*_*animal*_ = 30). The trimming activities for the sponge followed three patterns (Fig. 4C). They cut off (1) all four corners of a sponge, (2) one corner of the sponges elliptically, or (3) two corners of the sponge linearly. The crabs trimmed the white area (Fig. 4C) and started excavating a cavity (Fig. 4C Twenty-three crabs skipped the trimming behavior in 33 trials.

**Figure 4.**
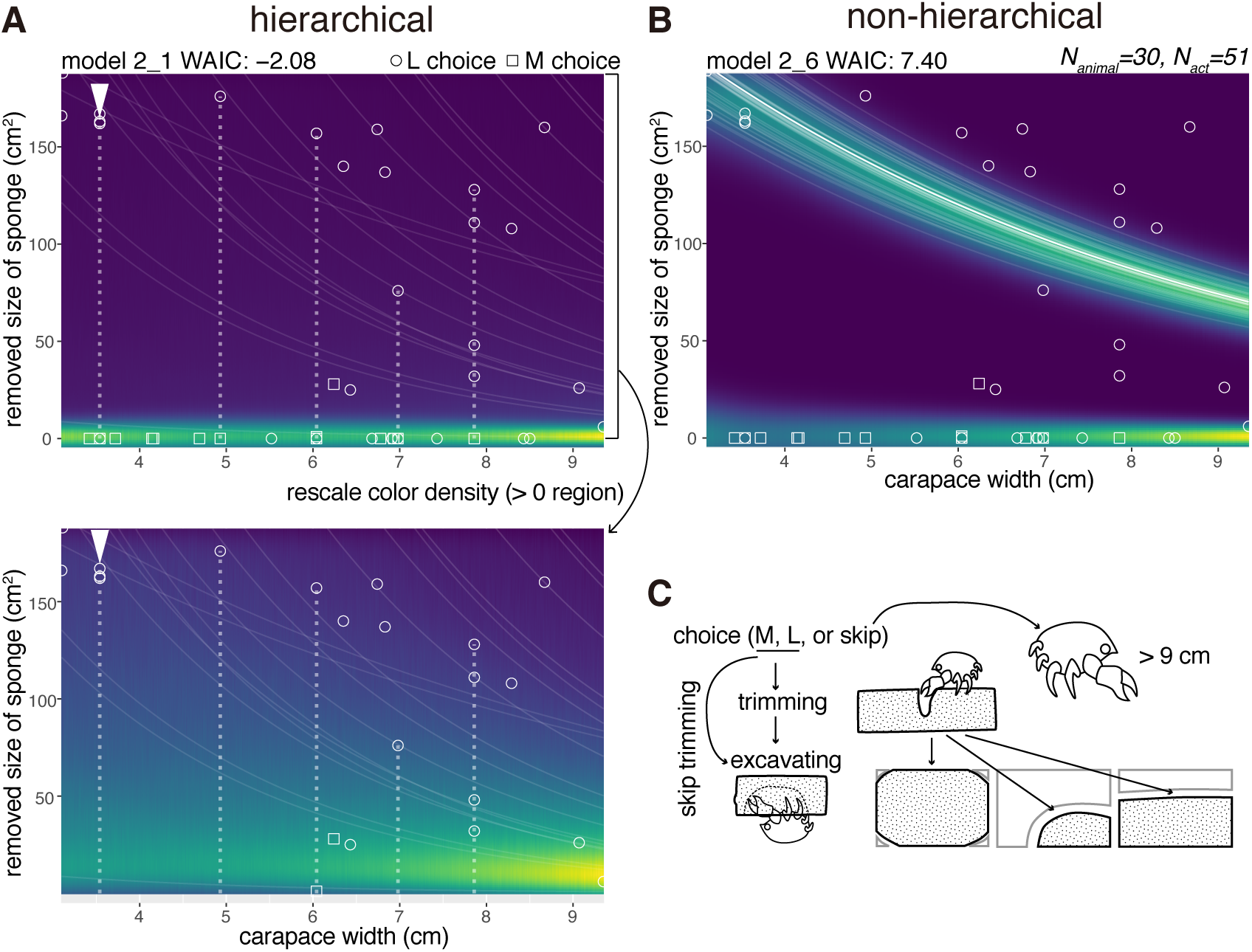
Trimming. (A) Upper plot: the predictive distribution on the hierarchical model 2_1. The white dotted lines connect the data points from the same individual. Lower plot: the predictive distribution visualized by re-scaling the color density of the expanded area in the upper plot except for the zero in the y-axis. (B) Predictive distribution on the non-hierarchical model 2_6. (C) Outline of the trimming process from a choice of a sponge (animals larger than about 9 cm skipped the whole behavior); some of the animals skipped the trimming behavior and went directly to cavity making. The drawing is by the authors.

After trimming or skipping, they started excavating. For the behavioral act of trimming, a non-zero data point indicating a trimmed size of the sponge was recorded (Fig. 4A,B). The size decreased with the increase of the carapace width. If a crab skipped trimming, a data point was recorded at zero (Fig. 4A), meaning no trimming. Almost all the crabs that chose the M size sponges decided not to trim the sponge except for one individual. Meanwhile, they trimmed less amounts of the sponges relative to the increase of their body sizes when they chose the L size sponges.

The WAIC of the hierarchical model 2 1 was −2.08 and that of comparable non-hierarchical model 2_6 was 7.40 (9.48 nat/sample in difference, Fig. 4D, Table 1), indicating that the hierarchical model is significantly better than the non-hierarchical one.

### Cavity size

Six crabs just cut the sponge and did not excavate the sponge. We modeled the size of a cavity in a cap (*N*_*animal*_ = 30) as a random variable taken from the gamma distribution with the log link function (Fig. 5). The size increased with the carapace width, and the model considering individual behavioral types performed best (Table 1). The WAIC of the hierarchical model 3_1 is slightly smaller than that of the comparable non-hierarchical model 3_2 (0.08 nat/sample in the difference; Fig. 5A,B, Table 1). The individual with the arrowhead (Fig. 5A) made relatively large cap cavities, indicating large inter-individual differences. As expected, larger crabs made larger cavities. The difference of the WAIC was about 0.1 (Fig. 5B). The predictability improvement is relatively small against that of sponge choice, suggesting that the individual behavioral type would have a lower effect in the determination of cavity size.

**Figure 5.**
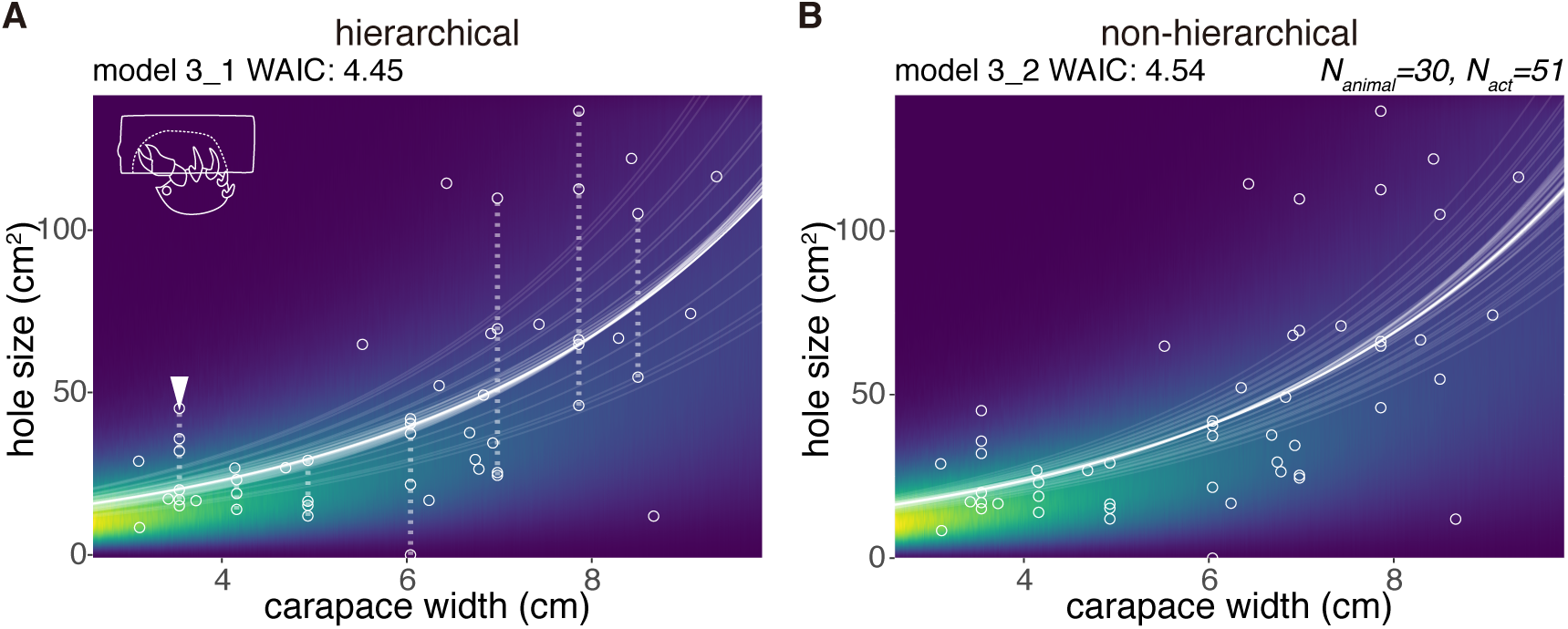
Excavated cavity in a cap. (A) Predictive distribution of a cavity size on the model 3_1. The white points connected by dotted lines are from the same individual. (B) Predictive distribution on the model 3_1. The drawing is by the authors.

### Latency

We modeled the latency for cap making (from the choice of sponge to carrying) by 32 crabs as a random variable taken from the zero-inflated distribution (Fig. 6). No obvious relation was found between the carapace width and the latency, and the number of crabs that had carried the cap by the next day. However, the hierarchical model 4_1 outperformed the non-hierarchical model 4_2 (WAIC values, 1.10 and 1.28 respectively, thus 0.18 nat/sample in the difference).

**Figure 6.**
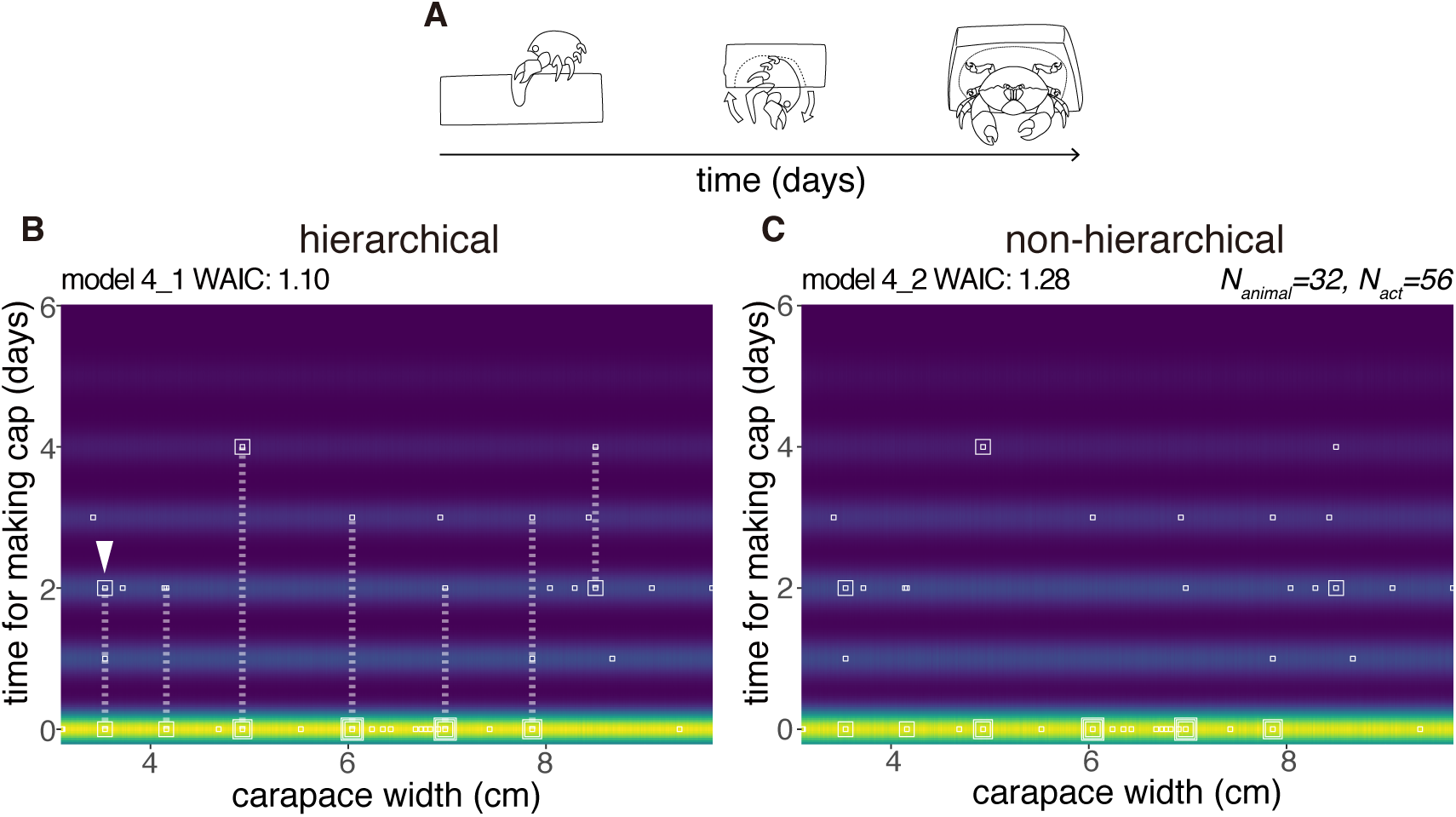
Latency to produce a cap. (A) Outline of cap making until carrying. (B) Predictive distribution of the latency on the model 4_1. Points from the same individual are connected by dotted lines. (C) Predictive distribution on the model 4_2. The drawing is by the authors.

## DISCUSSION

First, we provide an explicit rationale and mathematical basis of our statistical approach for the problem of quantification of the behavioral variability within and among individuals. Second, we discuss the empirical data of the sponge crab through the lens of our statistical framework.

### Statistical modeling from the predictive point of view

First, as a general theory, we state that the model construction for prediction is appropriate in our case because of the difference between prediction and discovery of true distribution. Second, from a mathematical point of view, we explain the maximum likelihood (ML) estimation and Bayesian inference, which are typical methods of statistical model construction, and discuss the validity of Bayesian inference and WAIC. Lastly, we consider the novelty of our statistical modeling in terms of the type of prediction and the difference between conditioned-level WAIC and marginal-level WAIC. Statistical models are broadly divided into those for prediction and those for discovering the true distribution. The two situations are as follows; (a) when there is no distribution that generates data in the finite set of models under consideration, (b) when there is a true distribution generating data in the finite set of models considered. Each situation is formulated as follows. Let *n* be the sample size. In case (a), the model is constructed by minimizing the generalization loss *G*_*n*_:

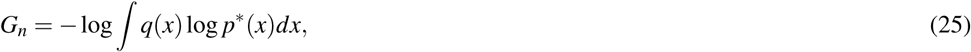

where *q*(*x*) is the true distribution of the data and *p*^∗^(*x*) is the predictive distribution. Predictive distribution is a probability distribution of the unknown new data based on the training data. For instance, the predictive distribution by ML estimation 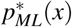 is defined by

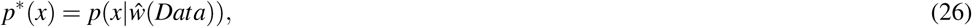

the likelihood model *p*(*x*|*w*) whose parameter *w* is plugged in the ML estimator (MLE) *ŵ*(*Data*). Bayesian predictive distribution is defined by the expectation of *p*(*x*|*w*) overall a posterior distribution *p*_*posterior*_(*w*|*Data*):

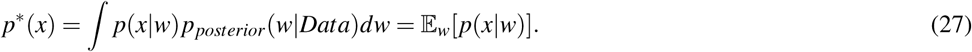

The generalization loss *G*_*n*_ is decomposed into two terms as below. The first term is independent from the model and the second one is the difference from the true distribution and the predictive one, called Kullback-Leibler divergence:

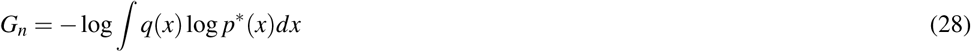

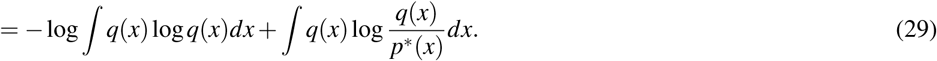

Thus, the generalization loss quantifies how far a predicted distribution is from the true distribution. In the case (b), the model is constructed by minimizing the negative log marginal likelihood *F*_*n*_:

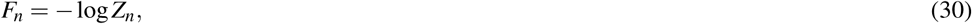

where *p*_*likelihood*_ (*Data*|*w*) is a likelihood, *φ*(*w*) is a prior distribution, and *Z*_*n*_ = *∫ p*_*likelihood*_ (*Data*|*w*)*φ*(*w*)*dw* is the marginal likelihood. Note that *Z*_*n*_ is equal to the normalizing constant of the posterior distribution since

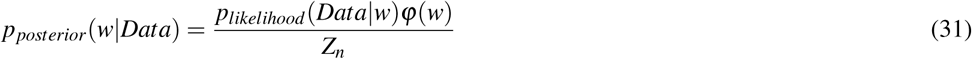

holds by the definition of conditional probability. In statistical mechanics, *F*_*n*_ is called ‘free energy’. Because of *Data* = (*x*_1_, …, *x*_*n*_), the true distribution of a data set is 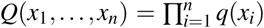. For simplicity, we put *x*^*n*^ = (*x*_1_, …, *x*_*n*_). We rewrite the marginal likelihood *Z*_*n*_ = *Z*(*x*^*n*^) to emphasize that it is a probability distribution of a data set. We consider the expectation of *F*_*n*_ overall *Q*(*x*^*n*^). Just like the generalization loss, it is decomposed into the model independent term and the Kullback-Leibler divergence from *Q*(*x*^*n*^) to *Z*(*x*^*n*^):

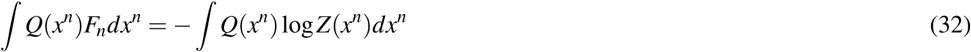

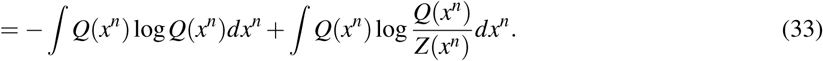

Thus, the expected free energy quantifies how different the marginal likelihood is from the true distribution. By definition, *Z*(*x*^*n*^) is the distribution of a data set, constructed by the model. In a practical application, a data set is only an obtained set, hence *F*_*n*_ is to be minimized.

For the above formulation, the following facts are known in statistics. In case (a), the ‘effective method’ would be a model selection method which can select the model while minimizing the generalization loss if *n* → ∞. Obviously, minimizing *G*_*n*_ is the ‘effective method’. Moreover, minimizing WAIC is also ‘effective’.

On the other hand, in case (b), a model selection method that can select the true distribution if *n* → ∞, is called ‘consistent method’. Although statistical modeling in case (b) is formulated by minimizing the expected free energy *∫ Q*(*x*^*n*^)*F*_*n*_*dx*^*n*^, minimizing the free energy *F*_*n*_ is ‘consistent’.

Our statistical analysis considers (a). We argue that no distribution generates data in the finite set of models, because our models are descriptive; they are not mechanistic models which represent individual behaviors of the crabs. Hence, it is appropriate to construct predictive models.

Next, we explain estimation methods. ML estimation and Bayesian inference are the typical methods used. However, they can be understood in a unified framework.

An analyst arbitrarily designs the simultaneous distribution of a pair (*x, w*)

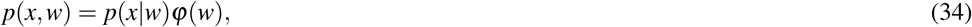

where *x* is an observable variable and *w* is a latent variable. In the ML estimation, the existence of the MLE *ŵ* is assumed and *φ*(*w*) is set to *δ*(*w* − *ŵ*) whose realization is limited to the MLE *ŵ*. As a formality, this can be interpreted as the parameter *w*, which is the MLE *ŵ*.

Let *w* be a real number. For an arbitrary real number *a*, the function *δ*(*w*−*a*) satisfies

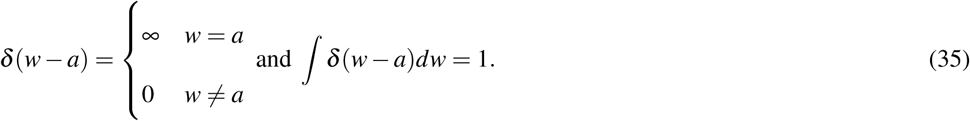

Thus, it is clear that *p*(*Data*|*w*)*δ*(*w*− *ŵ*)*/Z*_*n*_ = *δ*(*w*− *ŵ*) holds.

In Bayesian inference, *φ*(*w*) is a prior distribution. ML estimation can be described in the same way as the Bayesian inference; ML estimation is the case when the prior distribution is fixed to *δ*(*w* − *ŵ*). Accordingly, ML estimation and Bayesian inference can be understood in a unified way for the parameter estimation. The method to be used depends on the purpose of statistical modeling.

In the construction of a predictive distribution, the following theorem has been proven. Let 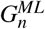 be the generalization loss of ML estimation and 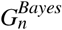 be the one of Bayesian inference. The overall expectation symbol across the dataset is denoted by 𝔼[*·*] = *∫ Q*(*x*^*n*^)[*·*]*dx*^*n*^. There are the constants *µ, λ* (0 < *λ* < *µ*), which are dependent on a model and the true distribution such that

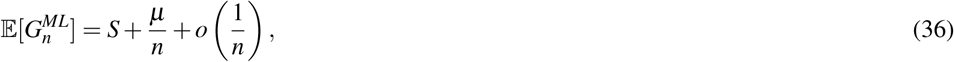

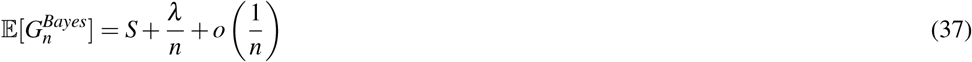

hold, where *S* = − *∫ q*(*x*) log *q*(*x*)*dx* is the entropy. Especially, when a prior distribution in Bayesian inference is strictly positive and bounded (0 < *φ*(*w*) < ∞),

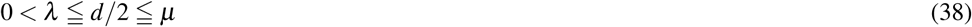

holds, where *d* is the dimension of the parameter. If the likelihood and the posterior distribution can be approximated by a normal distribution, the equal signs hold. In this study, we use hierarchical models. For them, the likelihood and posterior distribution cannot be approximated by *any* normal distribution. Therefore, the Bayesian inference can make the generalization loss smaller than that of ML estimation, i.e., it is appropriate for constructing a predictive model:

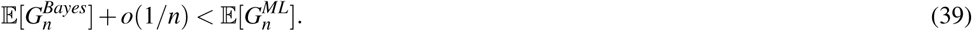

Third, we discuss the evaluation criterion used in our analysis. We consider the case (a) as appropriate, so we are to minimize the generalization loss. In our case, the model selection should be ‘effective’, thus neither using ML estimation nor the coefficient of determination *R*^2^ is appropriate. However, unfortunately, the generalization loss cannot be computed since the true distribution *q*(*x*) is unknown. When a model is set in which the likelihood can be approximated by a normal distribution, the Akaike information criterion (AIC) can estimate the generalization loss 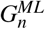 with the theoretical proof (Akaike, 1974). Moreover, minimizing AIC is the ‘effective method’ if the above assumption is satisfied. Hence, let *A*_*n*_ be AIC

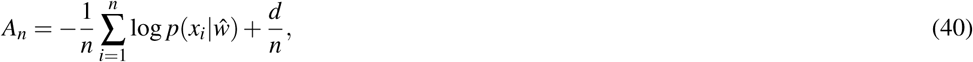

we have

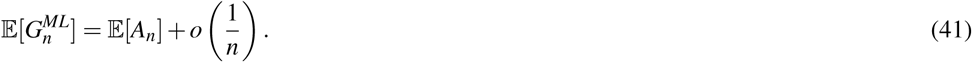

Note that we write the AIC in the scale of the generalization loss, not the deviance scale (2*nA*_*n*_). As mentioned above, our model set includes a hierarchical model, thus it is not appropriate to minimize AIC in the ML estimation. We chose Bayesian inference in order to decrease the generalization loss 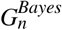. In addition, there are other advantages of the Bayesian inference. The widely applicable information criterion (WAIC) Watanabe (2010a) can estimate 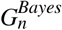 with mathematical proof even if the likelihood and the posterior cannot be approximated by any normal distribution. Moreover, WAIC is “effective”. Let *W*_*n*_ be WAIC

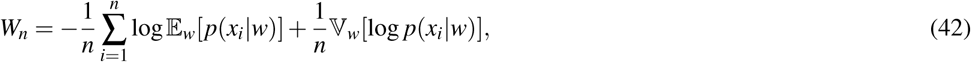

we have

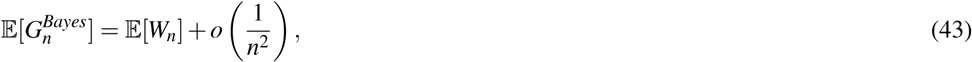

without the assumption of normality for the posterior distribution Watanabe (2010a).

Another model evaluation criterion is well-known: widely applicable Bayesian information criterion (WBIC) Watanabe (2013). Let 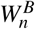 be WBIC. WBIC approximates the free energy: 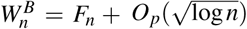 Watanabe (2013). It is useful for statistical modeling of case (b). However, in our case, we do not suggest that the mechanistic model is included in the considered model set; instead, we perform a statistical modeling of case (a). Therefore, we concluded that the model evaluation using WAIC was more appropriate than using WBIC. Indeed, WAIC is useful for real data even if we are limited to behavioral data (Wakita et al., 2020; Barrett et al., 2017).

Lastly, we discuss the difference between conditioned-level WAIC and marginal-level WAIC. Although WAIC is beginning to be used for evaluating models with empirical data, we should be careful to compute the value of a hierarchical model. Watanabe (2018) introduces two different definitions of WAIC depending on two different predictions. The often-used definition of WAIC for a hierarchical model is the first case in the book:

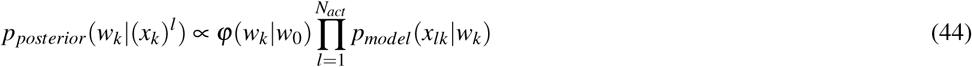

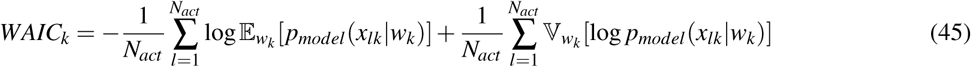

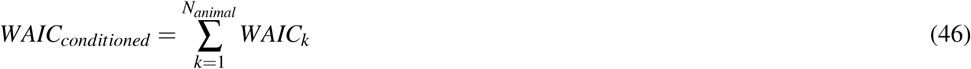

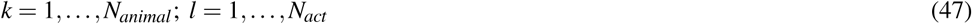

where 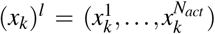 represents all given data for an individual. Note that the number of behavioral acts of the *k*-th animal is the same (balanced) for simplicity here (The number is unbalanced in our data). It should be noted that the statistical model *p*_*model*_ (*x*_*lk*_|*w*_*k*_) is conditioned upon the *w*_*k*_ assigned to each individual. In other words, in prediction, this model focuses on new acts of the already obtained individuals, whereas our focus is usually on a new act of another individual in order to compare models. In many cases, when studying ‘animal personality’, we are not usually interested in how our models specifically explain the sampled animals ‘personalities. Instead, we are interested in the distribution of a focused species. This is the reason why we did not use this conditioned-level WAIC. However, it is generally more convenient to use conditional-level likelihood within most Bayesian software, with the unfortunate consequence that conditional-level WAIC is often used. For example, Mitchell et al. (2016) uses conditioned-level WAIC to compare a hierarchical model with a non-hierarchical model. Furthermore, the use of the measure *R*^2^ for the evaluation of the model in terms of ‘variance explained’ is based on how we can minimize the variation in the sample obtained, and not focused on the prediction of the true distribution (Nakagawa and Schielzeth, 2010).

In summary, we took ‘a’ best approach from the predictive point of view and explored ‘a’ best model rather than a ‘correct’ model, because it is a natural assumption that the distribution we build would never be correct in the empirical modeling of a natural behavior of the sponge crab.

### Making cost and size choice: why did the crabs skip carrying the sponge?

The crabs in our experiments that did not carry caps were larger than those that carried caps. One possibility for the reason would be that when they grow up to some extent, the predators might avoid the crab and the relative energetic cost to make caps might increase. We speculate that this might be a reason why the large crabs did not make nor carry the caps.

Another possibility is that the sponges used in this experiment were smaller than those of the necessary size for the crabs. Dembowska (Dembowska, 1926) reported that the proportion of caps that fit the size of *D. personata* tended to decrease with the increase of the size of the crabs, and we considered that the decision to skip carrying the caps was because there were few sponges that fit the large crabs. Similarly, the large crabs that skipped cap making and carrying, would carry a cap if a sponge size would be larger than the L size sponge. In addition, no individuals carried the S sponge, because it was too small for all of the crabs to carry. It is likely that a younger and smaller crab than those used in this experiment would carry the S sponge. However, we can not exclude the possibility that the variations of the morphology of dactyls or the molt stage affect the behavior.

### Conclusion

We focused on the body extending behavior of the sponge crab, since the sponge crabs seemed to rely on the behavioral plasticity to make the living or non-living materials suitable to the animal body. Another crustacean that exhibits the body extending behavior is the hermit crab that is well known to prefer specific shells (Bertness, 1980; Hazlett, 1981; Wilber, 1990). Although hermit crabs cannot modify the shells by themselves, they are suggested to recognize and learn the shape of extended shells and the surrounding terrain (Sonoda et al., 2012). Therefore, the hermit crabs also might have behavioral types.

McLay (1983) showed the relationship of the body and cap size of the *Cryptodromia hilgendorfi* using a log link function and Gaussian distribution. As shown in the crab, we conditioned the variables on the carapace width of the crab *Lauridromia dehaani*. To consider ‘animal personality’, it is important to appropriately condition out the variables assumed to have much information about the behavioral variable. The body size is presumed to be an influential variable for the cap making behavior. Therefore, we conditioned all behavioral variables upon body size and found that the predictability improved by adding an assumption that all behavioral aspects pertained to the ‘individual. ‘The improvement was larger in the sponge choice than in the cavity size. Because the cavity size was determined by repeated excavation and body rotation, the crab might have used the carapace as a ‘measure’. However, in the choice task, the information processing to measure an appropriate size would be less dependent on the measure. We speculate that this makes a room for the emerging individual behavioral types dependent on behavioral plasticity that is unique to an individual.

## ACKNOWLEDGMENTS

We wish to thank Shirahama aquarium for allowing us to use the aquarium tanks, and Sakai fishing port authorities for providing the crabs. We wish to acknowledge Dr. Michael Rosario for his advice in improving the manuscript. We would like to thank Editage (www.editage.com) for English language editing.

## REFERENCES

Akaike, H. (1974). A new look at the statistical model identification. IEEE Transactions on Automatic Control, pages 716–723.

Akaike, H. (1980). Likelihood and the Bayes procedure. In J. M. Bernardo, M. H. De Groot, D. V. L. and Smith, A. F. M., editors, Bayesian Statistics, pages 143–166. Valencia: University Press.

Barrett, B. J., McElreath, R. L., and Perry, S. E. (2017). Pay-off-biased social learning underlies the diffusion of novel extractive foraging traditions in a wild primate. Proceedings of the Royal Society B: Biological Sciences, 284(1856):20170358.

Bedini, R., Canali, M. G., and Bedini, A. (2003). Use of camouflaging materials in some brachyuran crabs of the mediterranean infralittoral zone. Cahiers de Biologie Marine, 44(4):375–383.

Bertness, M. D. (1980). Shell preference and utilization patterns in littoral hermit crabs of the bay of panama. Journal of Experimental Marine Biology and Ecology, 48(1):1–16.

Dembowska, W. S. (1926). Study on the habits of the crab *Dromia vulgaris* ME. The Biological Bulletin, 50(2):163–178.

Gelman, A., Rubin, D. B., et al. (1992). Inference from iterative simulation using multiple sequences. Statistical science, 7(4):457–472.

Guinot, D. and Wicksten, M. K. (2015). Camouflage: carrying behaviour, decoration behaviour, and other modalities of concealment in brachyura. In Treatise on Zoology-Anatomy, Taxonomy, Biology. The Crustacea, Volume 9 Part C (2 vols), pages 583–638. Brill.

Hazlett, B. A. (1981). The behavioral ecology of hermit crabs. Annual Review of Ecology and Systematics, 12(1):1–22.

Hunt, G. R. (1996). Manufacture and use of hook-tools by new caledonian crows. Nature, 379(6562):249.

Kirschner, M. and Gerhart, J. (1998). Evolvability. Proceedings of the National Academy of Sciences, 95(15):8420–8427.

Matsui, H. and Izawa, E.-I. (2017). Flexible motor adjustment of pecking with an artificially extended bill in crows but not in pigeons. Royal Society Open Science, 4(2):160796.

McLay, C. L. (1983). Dispersal and use of sponges and ascidians as camouflage by *Cryptodromia hilgendorfi* (Brachyura: Dromiacea). Marine Biology, 76(1):17–32.

Millar, R. B. (2018). Conditional vs marginal estimation of the predictive loss of hierarchical models using WAIC and cross-validation. Statistics and Computing, 28(2):375–385.

Mitchell, D. J., Fanson, B. G., Beckmann, C., and Biro, P. A. (2016). Towards powerful experimental and statistical approaches to study intraindividual variability in labile traits. Royal Society open science, 3(10):160352.

Nakagawa, S. and Schielzeth, H. (2010). Repeatability for gaussian and non-gaussian data: a practical guide for biologists. Biological Reviews, 85(4):935–956.

Niemelä, P. T. and Dingemanse, N. J. (2018). On the usage of single measurements in behavioural ecology research on individual differences. Animal behaviour, 145:99–105.

R Core Team (2018). R: A Language and Environment for Statistical Computing. R Foundation for Statistical Computing, Vienna, Austria.

Reinhart, A. (2015). Statistics done wrong: The woefully complete guide. No starch press.

Sakamoto, Y., Ishiguro, M., and Kitagawa, G. (1986). Akaike information criterion statistics. Dordrecht, The Netherlands: D. Reidel, 81.

Sih, A., Bell, A., and Johnson, J. C. (2004). Behavioral syndromes: an ecological and evolutionary overview. Trends in ecology & evolution, 19(7):372–378.

Sonoda, K., Asakura, A., Minoura, M., Elwood, R. W., and Gunji, Y.-P. (2012). Hermit crabs perceive the extent of their virtual bodies. Biology letters, 8(4):495–497.

Stan Development Team (2018). Stan Modeling Language Users Guide and Reference Manual Version 2.18.0.

Wakita, D., Kagaya, K., and Aonuma, H. (2020). A general model of locomotion of brittle stars with a variable number of arms. Journal of the Royal Society Interface, 17(162):20190374.

Wang, L., Brodbeck, L., and Iida, F. (2014). Mechanics and energetics in tool manufacture and use: a synthetic approach. Journal of the Royal Society Interface, 11(100):20140827.

Watanabe, S. (2005). Algebraic geometry of singular learning machines and symmetry of generalization and training errors. Neurocomputing, 67:198–213.

Watanabe, S. (2010a). Asymptotic equivalence of Bayes cross validation and widely applicable information criterion in singular learning theory. Journal of Machine Learning Research, 11(Dec):3571–3594.

Watanabe, S. (2010b). Equations of states in singular statistical estimation. Neural Networks, 23(1):20–34.

Watanabe, S. (2013). A widely applicable Bayesian information criterion. Journal of Machine Learning Research, 14(Mar):867–897.

Watanabe, S. (2018). Mathematical theory of Bayesian statistics. Florida: CRC Press.

Wilber, T. (1990). Influence of size, species and damage on shell selection by the hermit crab *Pagurus longicarpus*. Marine Biology, 104(1):31–39.

Zuur, A., Ieno, E. N., Walker, N., Saveliev, A. A., and Smith, G. M. (2009). Mixed effects models and extensions in ecology with R. New York, USA: Springer.

