## Supplementary figures and images for "Individual behavioral type captured by a Bayesian model comparison of cap making by sponge crabs"

### supplementary figure 1

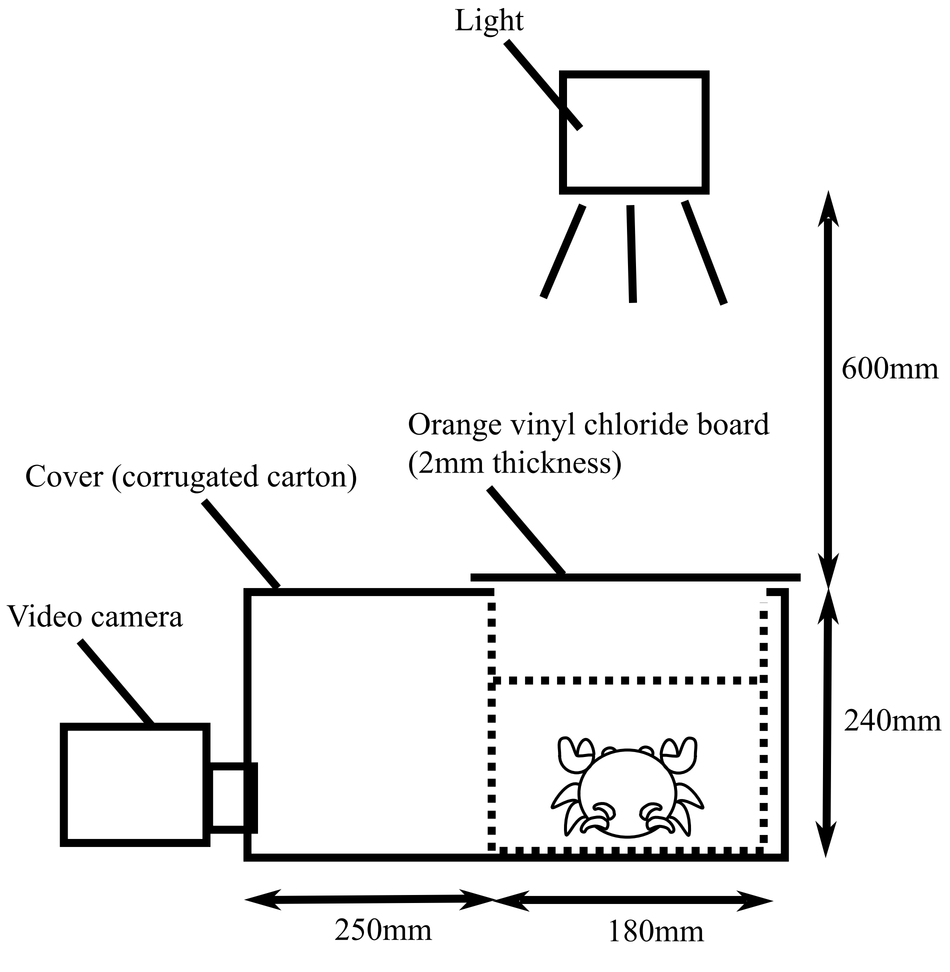
